# Metrics for Evaluating Biological AI Model Prediction Accuracy at the Data-Substrate Level

**DOI:** 10.64898/2026.06.14.732129

**Authors:** M. Antony Ewing

## Abstract

**Summary:** Reports in the biological literature disagree on whether a given model can predict a biological outcome from a given data sample — one study finding a model capable, another, on the same kind of data, finding it is not. This is particularly a challenge in relation to LLMs–where the models are large and opaque, with weights and training data inaccessible. Such disagreements cannot be settled by directly inspecting the model. To address this challenge, we consider an alternative approach: assessing whether the data sample is adequate to support the prediction asserted. For a given dataset, its substrate — the underlying structure of the data — determines what any model can recover, independent of architecture or capacity. At the same time, predicting the present state of a biological process and predicting the direction of its future change are different tasks; the second is supportable among AI models only where the data encode direction as determinable from the state — a property we call encoding — and is unsupportable where the same observed state precedes change in opposite directions — a property we call non-identifiability, in the informational rather than the statistical sense. We introduce two generic metrics, Predictive Blindness Risk (PBR) and Prediction Indeterminacy Measure (PIM), that evaluate a data substrate for predictive accuracy directly — without access to model weights, architecture, or training data — and locate the regions of a data substrate where a predictive claim can be supported and where it cannot. Using human biological subjects, we employ the Yale Brain Metastases Longitudinal Data (1,430 human subjects; 11,892 MRI studies; four sequences) and show that direction of change was non-identifiable across regions encompassing the majority of transitions; a nonlinear AI model gained essentially nothing over majority-direction prediction there while recovering direction near-perfectly where the state encoded it; and model accuracy tracked data-substrate resolvability continuously (Spearman ρ = −0.95 to −1.00). The metrics adjudicate, before any model is trusted and from the data alone, where claims of predictive accuracy — of state, or of the law of change — can be supported.

**Background:** Model prediction of a biological outcome from a given data sample is often highly contested — particularly in predicting biological change. The literature reports conflicting capability for the same kind of data and when complex models or opaque LLMs are used, the disagreement cannot be resolved by inspecting the model or its benchmark training data. Such disagreement has two sources — instability of the model, and indeterminacy of the data — of which only the first has been studied. We address the second by relocating the question from the model to the data substrate.

**Methods:** We analysed human biological data taken from the Yale Brain Metastases Longitudinal Data (1,430 human subjects; 11,892 MRI studies; four sequences) using transitions between consecutive scans. An observed-measure proxy grouped subjects into matched observed states. We defined two data-substrate metrics — Predictive Blindness Risk (PBR), the minority-direction fraction at a matched state, and Prediction Indeterminacy Measure (PIM), the separation between admissible trajectories at a state — and evaluated state-only prediction with subject-level-split logistic regression and a multilayer perceptron against a majority-direction baseline. At bin level across all eight bins per sequence, we computed the Spearman correlation between PBR and held-out model accuracy. Formal results established when a single-valued predictor cannot recover direction at a shared state (Appendix); a controlled simulation with a known governing law served as comparison.

**Results:** Across sequences, 6–7 of 8 matched-state bins contained transitions in both directions, encompassing 75–88% of transitions; weighted PBR ranged 0.30–0.35 (mean 0.33, SD 0.01). In these non-identifiable bins the nonlinear model did not exceed majority-direction prediction (model−majority −0.014 to +0.001 across sequences and splits); in the complementary encoding bins (12–25% of transitions) accuracy reached 0.958–1.000. Across all bins, model directional accuracy declined monotonically with PBR (Spearman *ρ* = −0.952 FLAIR, −0.976 T1 post-contrast, −1.000 T1 pre-contrast, −0.976 T2; P ≤ 3.3×10^−5^ in three sequences, 2.6×10^−4^ in FLAIR), establishing a continuous gradient of data-substrate resolvability rather than a binary partition. PBR was stable across binning resolution (range 0.299–0.353; CV 5.4%). In the simulation the state-only model predicted the wrong direction for every minority-direction subject while retaining significant overall association (*ρ* = 0.46); theoretical PBR tracked empirical failure (*ρ* = 0.96).

**Conclusions:** State-only prediction recovers directional structure where the data substrate encodes it and cannot where the substrate is non-identifiable, with model accuracy varying continuously between the two as a near-deterministic function of data-substrate resolvability. The contested question of model predictive capability is therefore, in part, a prior question about the data: PBR and PIM answer it from the data alone, without access to model weights, architecture, or training data, and discriminate the regions of a data substrate where a claim of predictive accuracy — of state, or of the law of change — is and is not supportable.

## Introduction

Reports in the biological literature frequently disagree on whether a given model predicts a given quantity from a given data sample: one study finds a model capable of predicting a biological outcome, while another, on the same kind of data, finds it is not.^1–4^ The problem is particularly acute in comparing Large Language Models (LLMs): where the models are large and opaque — their weights and training data inaccessible to the parties evaluating them, as is increasingly the case for AI applied to biological processes. This disagreement cannot be settled by inspecting the model, because the only available evidence is its predictions, and predictions alone do not reveal whether a capability is genuine or whether a failure reflects the model or the data it was given.

Such disagreement has two sources, and only one is a property of the model. A model may be unstable — returning different outputs on repeated evaluation^3,4^ — which is a question of the model’s reliability and is studied as such. But a model may also be asked to predict something the data substrate does not determine. Where the data do not contain what is needed to recover the answer, different evaluations recover the available partial signal to differing degrees, and report conflicting capability for reasons that no amount of model refinement or re-testing will resolve. This second source is a property of the data substrate, not the model, and it is not currently measured.

This second challenge is most consequential for the prediction of biological *change*, as distinct from biological *state*. Identifying the state of a process at a point in time, and predicting the direction in which that state will change, are different tasks: the first is a reading of the present, the second a claim about a law governing the process. The first task may be written as predicting state y from observed state x in the data. The second task can be expressed as predicting Δy = f(x, *φ*), where x is the observed state and *φ* the determinants of its direction. Direction is recoverable from x only where *φ* is expressed in x (that is, x(*φ*), enabling the capture of *φ* from better sampling. In that case, predicting state and predicting change coincide, and a model that has captured the state predicts change as well. We refer to this situation as *encoding*, because the data encode direction as determinable from the state.

Where *φ* is not expressed in x, the same observed state precedes change in opposite directions, and no model — of any class, at any scale — can recover direction from x alone, even with further sampling. We refer to this latter property of the data as *non-identifiability* — not in the probabilistic or statistical sense of parameter or causal identifiability,^5,6^ but in the informational sense: direction is ambiguous and non-determinable from the state, because the information required to determine it is not present in the data, even if model parameters are identifiable in the traditional sense. A claim to predict change is supportable under encoding and not under non-identifiability, and which property a region of the data substrate exhibits is a fact about the data and the subject of this paper.

We present a set of metrics that evaluate a data substrate for predictive accuracy directly — without access to model weights, architecture, or training data — and that locate, within a data substrate, the regions where a predictive claim — whether of the state or of the future state direction — can be supported and the regions where it cannot. These do so by determining regions of encoding and non-identifiability, respectively. Predictive Blindness Risk (PBR) quantifies the fraction of cases at a matched observed state that follow the minority direction — the rate at which any state-only predictor would be directionally wrong under non-identifiability. Prediction Indeterminacy Measure (PIM) quantifies, at a given state, how far apart the admissible trajectories are, identifying states compatible with change of opposite sign. Both are computed from longitudinal observations alone, are reproducible across implementations, and apply to a model an evaluator cannot open. Their inverses prescribe the regions of adequate potential prediction.

We establish the conditions formally — the conditions under which a single-valued predictor provably cannot recover direction of change at a shared state (Proposition A0, Corollary A0.2, Theorem A1, Appendix) — and demonstrate the metrics on a longitudinal biological imaging data substrate in human biological subjects, the Yale Brain Metastases Longitudinal Data^7^: 1,430 human subjects, 11,892 MRI studies, four sequences. Across all four sequences, the observed state leaves the direction of change non-identifiable across the regions encompassing the majority of transitions; a nonlinear model gains essentially nothing over majority-direction prediction in those regions (−0.014 to +0.001 across sequences) while recovering direction near-perfectly (0.96–1.00) in the regions where the state encodes it; and model directional accuracy tracks the data-substrate metrics continuously, as a near-deterministic function of how much direction the state encodes (Spearman *ρ* = −0.95 to −1.00 across sequences). PBR and PIM index where in this division any region of the data substrate falls, from the data alone, before reliance on a model’s directional output is established.

## Methods

### Formal Framework

For an observed state x (here, a lesion-burden proxy derived from MRI) and latent determinants *φ* (biological factors not expressed in the observed state at the time of prediction), the trajectory change is given by Δy = f(x, *φ*), where f is the law governing the biological trajectory. The predictor h(x; *θ*) must return a single value at each input. When two contextual regimes *φ*_1_ and *φ*_2_ yield f(x, *φ*_1_)≄ f(x, *φ*_2_) at a shared observed state x, no single-valued function of x can recover both. This is a structural feature of the prediction task, not a limitation of model capacity or training-data volume, and it holds regardless of whether *φ* is measurable in principle.

Two measures quantify the severity of this structural limit. PBR(x) := sup{|f(x, *φ*_1_) − f(x, *φ*_2_)| : *φ*_1_, *φ*_2_ ∈ Φ} equals zero if and only if direction at x does not depend on context. PIM(x) := |f(x, *φ*_1_) − f(x, *φ*_2_)| / max{|f(x, *φ*_1_)|, |f(x, *φ*_2_)|}, with PIM > 1 indicating admissible trajectories of opposite sign. Proofs are in the Appendix. Empirically, we estimate both from observed branching structure in longitudinal data. The empirical operationalisation is an observable analogue that does not require specification of f or identification of *φ*.

### Data

The Yale Brain Metastases Longitudinal Data is hosted on The Cancer Imaging Archive (TCIA). It comprises 1,430 subjects across 11,892 MRI studies and four sequences: FLAIR (9,083 studies), T1 post-contrast (8,996), T2 (7,348), and T1 pre-contrast (8,384). The cohort reflects a predominantly adult population with brain metastases from lung and breast primary cancers, consistent with the epidemiology of adult brain metastasis. No clinical annotations or treatment labels are included. This study used publicly available, de-identified data; ethics approval was not required under 45 CFR 46.104(d)(4).

### Lesion Proxy and Transitions

From each MRI volume, we extracted a scalar lesion-burden proxy via Gaussian smoothing, robust z-score normalisation, bright voxel cluster identification, connected-component filtering, and integration of cluster volume and mean intensity. A multi-threshold fallback ensured proxy extraction from scans with diffuse signal. The proxy is an ordinal summary designed to group subjects into similar observed states; it is not a calibrated biological measurement. Branching is visible in raw longitudinal transitions prior to binning, independent of the specific proxy construction. For each subject with at least two scans of a given modality, consecutive-scan pairs within 180 days yielded transitions (proxy at scan t; Δy = proxy at scan t+1 minus proxy at scan t). A deadzone filter excluded the weakest 20% of |Δy| values to remove near-zero changes attributable to measurement noise.

### Empirical Estimation of PBR and PIM

Transitions were grouped into eight equipopulated bins of the lesion proxy. Within each bin, we computed the minority-branch fraction. A bin was classified as *branching* if the minority fraction exceeded 10%. Empirical PBR was estimated as the weighted average minority fraction across branched bins. The empirical operationalisation of PIM was 1 minus the accuracy of the state-only model in branched bins. The fidelity of these empirical estimators to their formal definitions is validated in the Appendix (§A7, §B), where the theoretical PBR correlated with observed sign-error rate at Spearman *ρ* = 0.96 in the Gompertz simulation; binning sensitivity is documented in §A7.

### Models and Evaluation

Two AI family models were trained on each modality to predict trajectory direction from the lesion proxy, selected for contrast: logistic regression and a multilayer perceptron (two hidden layers: 32, 16 units; ReLU; Adam; early stopping). Subject-level 70/30 splitting ensured no subject appeared in both sets. Each modality–model combination was evaluated across three independent random seeds. Reported values are means ± SD across splits. PBR stability was assessed across four bin configurations (6, 8, 10, 12), yielding 16 modality–bin combinations.

### Substrate-Resolvability Gradient Analysis

To test whether the data substrate measures index a continuous gradient of resolvability rather than a binary partition between branching and non-branching regions, we computed, for each modality and within each of the eight equipopulated bins, two quantities on the held-out test predictions: the minority-branch fraction (a bin-level measure of data substrate indeterminacy) and the test accuracy of the multilayer perceptron (a bin-level measure of model directional recovery). Spearman rank correlation between minority-branch fraction and model accuracy across the eight bins quantified the gradient per modality. Under the framework’s prediction, this correlation should be strongly negative — data substrate-indeterminate bins should bound model accuracy from above, and data substrate-resolvable bins should permit recovery near the ceiling.

### Gompertzian Comparison Reference

To provide a reference point in which the governing law is known exactly, we included a controlled simulation using Gompertzian dynamics^8^ with two contextual regimes (progression and remission) and overlapping observed states. In this idealised setting, theoretical PBR and PIM are computed directly from f(x, *φ*), and a state-only predictor recovers the prevalence-weighted answer in branched bins. The simulation serves only as a comparison reference for the Yale empirical results; full parameters and extended sensitivity analyses are reported in the companion preprint.^9^

## Results

### Dataset

After proxy extraction, 1,271 to 1,362 subjects per modality (of the 1,430 total) contributed 4,185 to 5,461 transitions after deadzone filtering (Table 1). Proxy extraction succeeded for over 99.9% of scans.

**Table 1.**
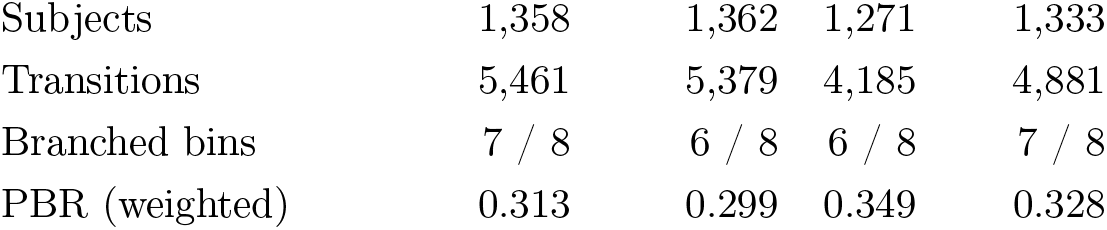
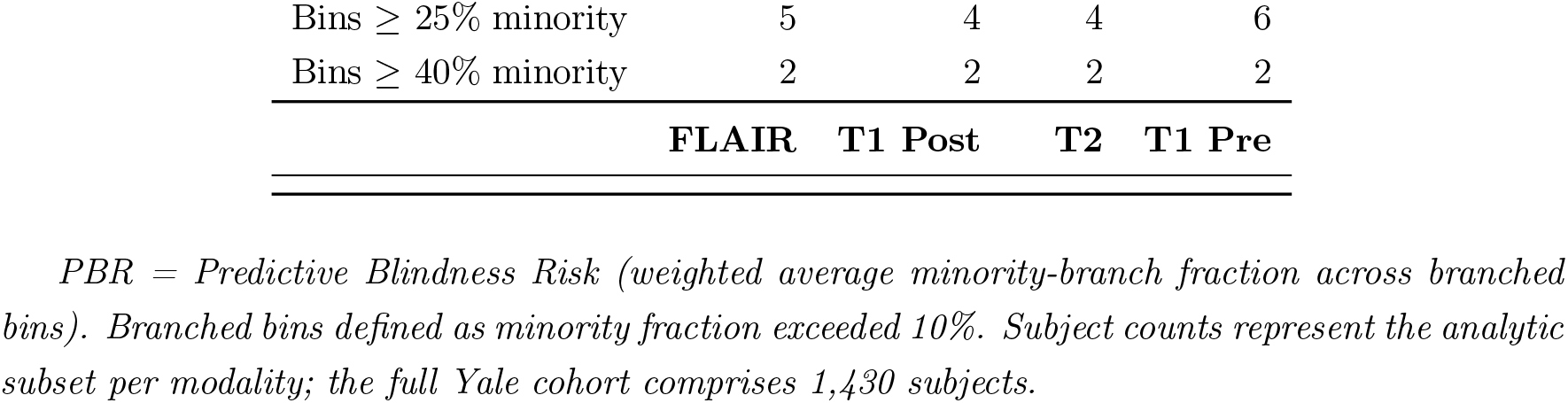
Non-identifiability of directional change across four MRI sequences in the Yale Brain Metastases Longitudinal Data.

|  |  |  |  |  |
| --- | --- | --- | --- | --- |
| Subjects | 1,358 | 1,362 | 1,271 | 1,333 |
| Transitions | 5,461 | 5,379 | 4,185 | 4,881 |
| Branched bins | 7 / 8 | 6 / 8 | 6 / 8 | 7 / 8 |
| PBR (weighted) | 0.313 | 0.299 | 0.349 | 0.328 |
| Bins ≥ 25% minority | 5 | 4 | 4 | 6 |
| Bins ≥ 40% minority | 2 | 2 | 2 | 2 |
|  | **FLAIR** | **T1 Post** | **T2** | **T1 Pre** |

*PBR = Predictive Blindness Risk (weighted average minority-branch fraction across branched bins). Branched bins defined as minority fraction exceeded 10%. Subject counts represent the analytic subset per modality; the full Yale cohort comprises 1,430 subjects.*

### Non-Identifiability of Direction Across Sequences

Matched-state bins contained transitions in both directions in every sequence (Figure 1). Of eight bins per modality, six to seven were branching, encompassing 75–88% of transitions. Weighted PBR ranged from 0.30 (T1 post-contrast) to 0.35 (T2); approximately one in three subjects at a given observed state followed the opposite trajectory from the majority (Table 1). Every modality contained bins with minority fractions exceeding 40%.

**Figure 1.**
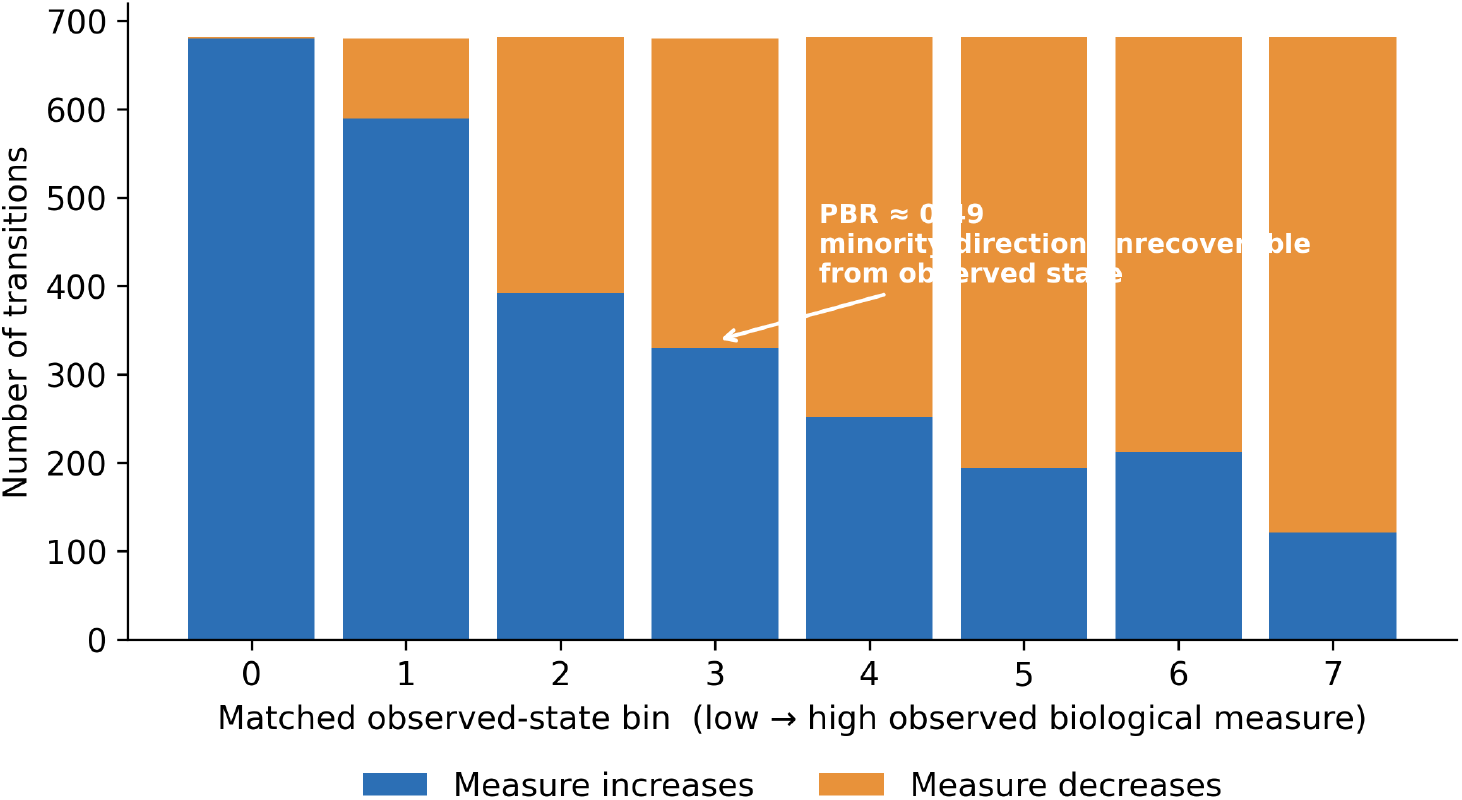
Non-identifiability of directional change within matched observed states (FLAIR). Transitions are grouped into bins of the observed biological measure (a lesion-burden signal from MRI), low to high. Within each matched-state bin, the measure subsequently increases for some human subjects and decreases for others. Under encoding (lowest bins) the direction is nearly uniform and recoverable from the state; as the observed state rises, the same state precedes change in opposite directions, and a state-only predictor is necessarily wrong for the minority share. That share is the Predictive Blindness Risk (PBR) for the bin (annotated). The three remaining sequences are shown in Supplementary Figure S2.

Across all four modalities, directional accuracy for a nonlinear neural network did not exceed majority-direction prediction in non-identifiable bins (Table 2, Figure 2). The MLP–majority gap ranged from −0.014 (T1 post-contrast) to +0.001 (T2, T1 pre-contrast) across three random splits, with positive gains not exceeding the within-split standard deviation. The measures therefore do not flag regions of simple-model underfitting; they flag regions in which increased nonlinear capacity does not resolve the directional task.

**Table 2.**
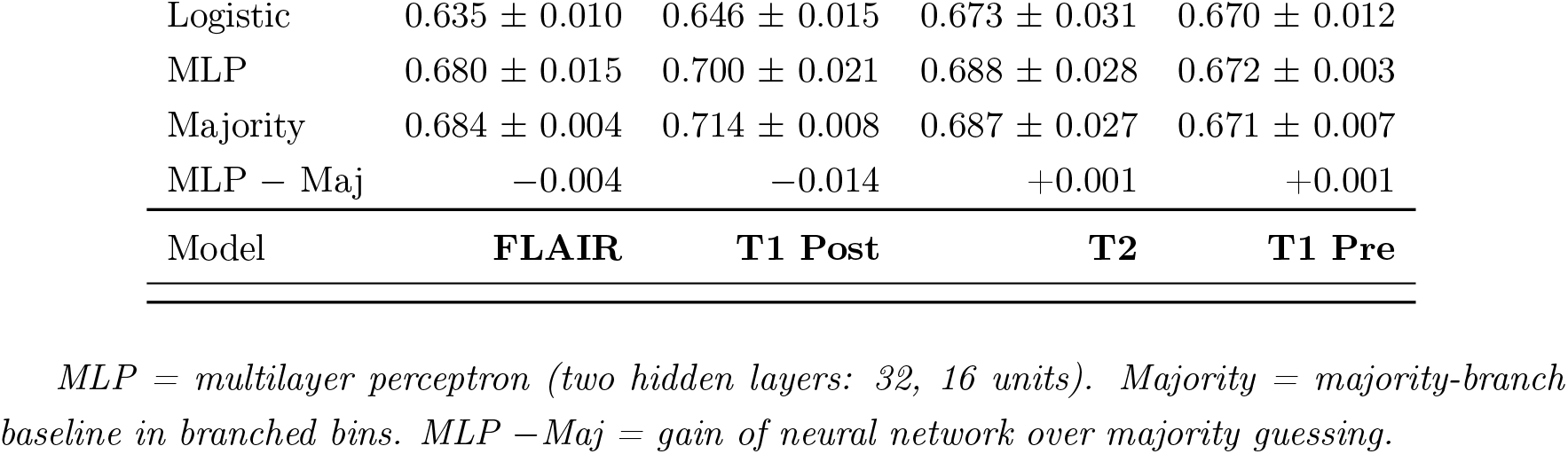
Directional accuracy in branched bins: logistic regression, neural network, and majority-branch baseline (mean ± SD, 3 subject-level splits).

|  |  |  |  |  |
| --- | --- | --- | --- | --- |
| Logistic | $0.635 \pm 0.010$ | $0.646 \pm 0.015$ | $0.673 \pm 0.031$ | $0.670 \pm 0.012$ |
| MLP | $0.680 \pm 0.015$ | $0.700 \pm 0.021$ | $0.688 \pm 0.028$ | $0.672 \pm 0.003$ |
| Majority | $0.684 \pm 0.004$ | $0.714 \pm 0.008$ | $0.687 \pm 0.027$ | $0.671 \pm 0.007$ |
| MLP – Maj | $-0.004$ | $-0.014$ | $+0.001$ | $+0.001$ |
| **Model** | **FLAIR** | **T1 Post** | **T2** | **T1 Pre** |

*MLP = multilayer perceptron (two hidden layers: 32, 16 units). Majority = majority-branch baseline in branched bins. MLP – Maj = gain of neural network over majority guessing.*

**Figure 2.**
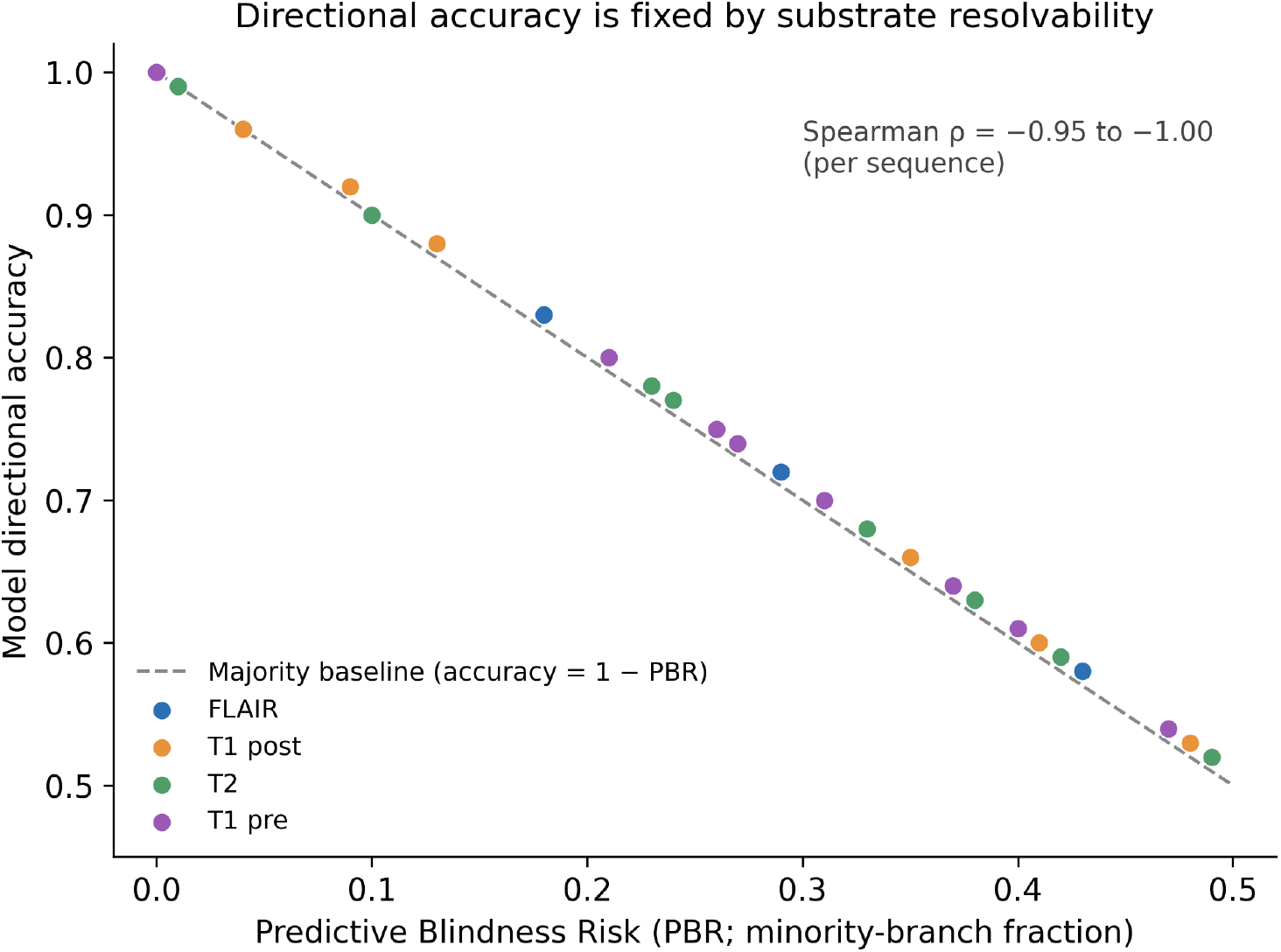
Model directional accuracy is fixed by data-substrate resolvability. Each point is one matched-state bin in one MRI sequence (four sequences, colour-coded). The horizontal axis is the bin’s Predictive Blindness Risk (PBR; minority-direction fraction); the vertical axis is held-out directional accuracy of a nonlinear model on that bin’s transitions. Points lie along the majority baseline (accuracy = 1 − PBR, dashed): as non-identifiability rises, attainable accuracy falls one-for-one, and added model capacity does not exceed the line. Per-sequence Spearman *ρ* between PBR and accuracy ranges from − 0.95 to − 1.00. Points are per-bin model accuracy, not per-subject; the dashed line is the analytic majority baseline, so the correspondence reflects a substrate limit a trained model does not surpass, not a fitted relationship.

### State-Only Prediction in Non-Branching Bins

In the complementary one to two non-branching bins per modality — the 12–25% of transitions where current state is already informative about direction — the same neural network achieved test accuracy of 1.000 in FLAIR (n=232), 0.958 in T1 post-contrast (n=623), 1.000 in T1 pre-contrast (n=173), and 0.958 in T2 (n=330). In these bins, where one trajectory direction dominates by construction, the majority-branch baseline is itself near-maximal (0.958–1.000); the model successfully recovers the directional structure the data substrate encodes rather than adding independent predictive value beyond it. The framework’s claim is symmetric: where the data substrate determines direction, the model recovers it; where the data substrate does not, the model cannot.

### Substrate-Resolvability Gradient Across All Bins

Across all eight bins per modality, model directional accuracy declined monotonically with the bin-level minority-direction fraction (Figure 2). Spearman rank correlation between minority fraction and model accuracy was −0.952 in FLAIR (P = 2.6×10^−4^), −0.976 in T1 post-contrast (P = 3.3×10^−5^), −1.000 in T1 pre-contrast (P < 1×10^−10^; perfect rank ordering across eight bins), and −0.976 in T2 (P = 3.3×10^−5^). The relationship is therefore not a binary partition into branching and non-branching regions but a near-deterministic gradient: data substrate indeterminacy at a bin progressively bounds the directional accuracy a model can attain on that bin’s transitions, with consistent functional form across four MRI sequences differing in physical basis. PBR and PIM index where on this gradient a given data substrate region falls.

### Encoding versus Non-Identifiability

The core empirical result can be summarised in a single contrast. Across modalities, the state-only model achieves overall test accuracy of 0.69–0.74 and AUC of 0.78–0.88 (Table 2) — performance levels that, by standard standard evaluation criteria, would support confidence in the model’s utility. Yet in the branched bins encompassing 75–88% of transitions, the same model’s directional accuracy (0.63–0.67) does not exceed the majority-branch baseline (0.65–0.71), meaning it has learned only which trajectory is more common, not which trajectory a given subject will follow. The model does not fail on a small edge case: it returns the majority-branch answer for every subject in matched-state bins where similar observed states were followed by opposite trajectories, and that answer is wrong approximately one in three times. Its response is also confident because the model does not represent the unresolved alternative — at a matched state where the training distribution contains both trajectories, the model assigns the majority trajectory to every subject as though the minority branch did not exist. Standard validation metrics — overall accuracy, AUC, Spearman *ρ* — aggregate across both determinate and branching regions and therefore conceal this directional failure precisely where it matters most. This is the state-to-state versus state-to-law gap: the model’s state-to-state capability is genuine, but its directional output in branching regions is structurally no better than guessing the prevalent direction.

### PBR Sensitivity

Across 16 modality–bin configurations, PBR ranged from 0.299 to 0.353 (mean 0.332, SD 0.014; coefficient of variation 4.3%). No configuration yielded PBR below 0.29 (main-text Figure 3).

**Figure 3.**
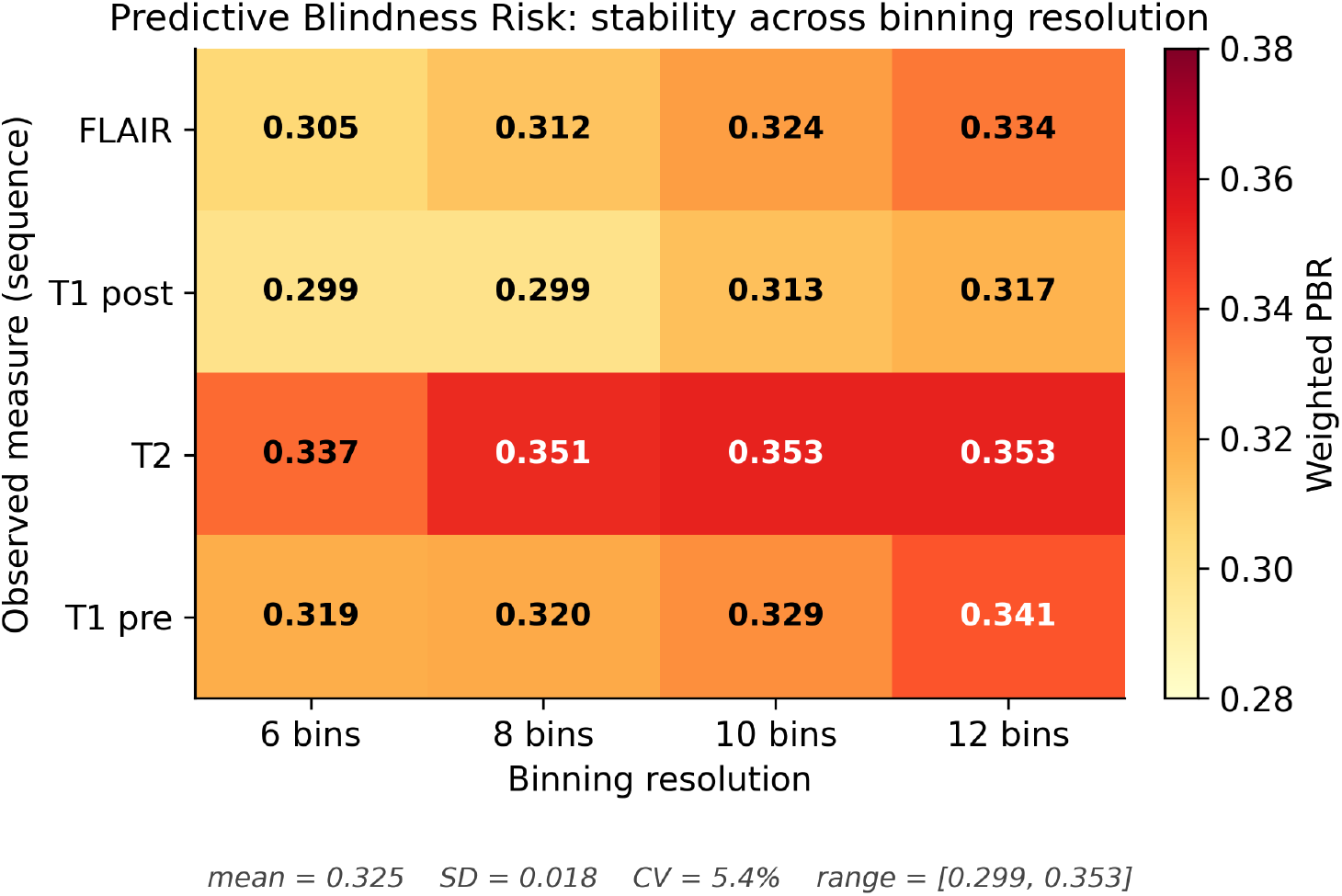
Predictive Blindness Risk is stable across binning resolution. Weighted PBR for each MRI sequence (rows) computed at four binning resolutions (columns). Values are tightly clustered (mean 0.325, SD 0.018, coefficient of variation 5.4%, range 0.299–0.353), so the measured non-identifiability is a property of the data substrate rather than an artefact of a particular binning choice.

#### Gompertzian Comparison Reference

Within the controlled simulation, theoretical PBR across the shared-state window ranged from 0.019 to 0.037 and PIM exceeded 1 throughout, with the state-only predictor converging on the prevalence-weighted branch in branching regions. The empirical PBR values from Yale (0.30–0.35; Table 1) lie well above the idealised simulation’s theoretical range, consistent with the larger and more heterogeneous determinant structure of an empirical biological data substrate relative to a two-regime controlled setting. Extended simulation details are reported in the companion preprint.^9^

## Discussion

Consider two subjects at the same observed state whose forward trajectories diverge (Figure 1). Their observations are indistinguishable because the determinant required to separate their trajectories is not present in the observed state at all — not because the model was poorly trained. A state-only model, asked to return a single directional answer for each, returns the same answer — the majority-branch trajectory — and that answer is wrong for whichever subject belongs to the minority. Standard validation does not detect this. A model may retain strong aggregate directional association while being directionally wrong for subjects for whom directional accuracy matters most. In the Yale data across four MRI sequences, the state-only model did not exceed majority-branch prediction in branching regions encompassing 75–88% of transitions. In a simulation designed to offer a base comparison, the state-only model preserved substantial association (*ρ* = 0.46) while predicting the wrong direction for every minority-branch subject.

The mechanism is identifiability, not model capacity: when the input does not determine trajectory direction — whether because direction of change is not encoded, forward-looking determinants are not yet expressed, or because the same observed state is compatible with divergent outcomes — additional architectural complexity cannot resolve the branching. The four-modality MLP experiment confirms this empirically.

The data substrate measures discriminate, rather than veto. In the same data and using the same models, directional accuracy reached 0.958–1.000 in non-branching bins and converged on the majority baseline in branching bins, with a near-deterministic monotone gradient between the two limits (Spearman *ρ* between −0.952 and −1.000 across the four MRI sequences). The data substrate measures therefore distinguish data substrate regions where a directional claim is supportable from regions where it is not, and index the magnitude of that support continuously rather than partitioning the data substrate into pass and fail. The framework’s contribution is this gradient, not a binary verdict. A reviewer may note that non-branching bins by construction have a high majority-branch baseline, so model accuracy and baseline accuracy coincide in those bins; this is correct, and is precisely what “data substrate-encoded direction” means operationally — the model successfully recovers the direction the data substrate encodes, and the gradient quantifies how that recovery degrades as the data substrate becomes less informative.

PBR and PIM differ from standard metrics in what they evaluate. AUC and accuracy ask how well a model performs on average. PBR asks whether the observed state determines trajectory direction at all. Where PBR is zero, conventional validation is sufficient. Where PBR is substantial, the observed state does not determine trajectory direction, and conventional metrics can certify a model that is directionally wrong for a large minority of subjects. A model achieving high aggregate performance may therefore still be structurally unreliable for the question it is assumed to answer: in which direction will this subject’s biological trajectory move? Distribution-shift monitoring addresses a related but distinct failure mode in which the input distribution itself changes after fitting; the structural limit described here can produce directional error even under stable distributions, and the two forms of post-hoc vigilance are complementary rather than substitutes.

The underlying result is a characterization, not merely a failure mode. State-only prediction can recover subject-specific trajectory direction if and only if the observed state encodes the determinant governing that direction (Corollary A0.2, Appendix). Where the observed state carries *φ* — so that Δy = f(x) holds — prediction from x alone is a well-posed problem and a model can solve it. Where it does not, no single-valued predictor can recover direction regardless of architecture, scale, or training-data volume; the task is not harder, it is ill-posed. This reframes a quantitative debate about model improvement into a qualitative question about the data substrate: does the observed state encode what the model is being asked to predict? PBR and PIM answer that question empirically, before reliance is established, without requiring access to model internals.

Operationally, PBR partitions the imaging-state space into regions where directional output can be trusted and regions where it cannot — and more precisely, indexes a continuous gradient between the two limits. In regions of low PBR, the observed state determines trajectory direction, model accuracy approaches the data substrate-encoded ceiling (0.96–1.00 in our data), and standard validation is sufficient. In regions of high PBR, the same observed state is compatible with divergent trajectories, model accuracy collapses to the majority-branch baseline, and a directional prediction should be treated as an indicator of the more common trajectory in that region rather than as a subject-specific claim; additional information that determines *φ* would be required before a directional claim is supportable. The objective is not to reject directional prediction wholesale but to identify where the observed state can resolve direction and, where it cannot, to make that boundary visible before the model’s directional output is relied upon. PBR and PIM make that partition before reliance is established and without requiring access to model weights.

Cross-modality convergence of PBR within 0.30–0.35 across four sequences with different physical bases — fluid-sensitive FLAIR, contrast-enhanced T1, T2-weighted, and non-enhanced T1 — is consistent with biological heterogeneity rather than measurement artifact. If branching were driven by sequence-specific noise, PBR would vary substantially across modalities; instead, it converges. The same convergence appears in the data substrate-resolvability gradient itself: the Spearman *ρ* between minority-branch fraction and model accuracy falls within −0.95 to −1.00 across all four modalities, indicating that the relation between data substrate resolvability and model directional accuracy is a property of the data substrate-claim relationship rather than of any particular MRI sequence. The branching is visible prior to binning and without model fitting. Moreover, the branching structure we document is visible without reference to hidden demographic confounders; it appears at matched imaging states regardless of covariate adjustment.

The present results are not an argument against using such models; they are an argument for bounded reliance — distinguishing tasks for which the observed state is sufficient from those for which direction of change depends on determinants the model does not receive or cannot uniquely resolve. PBR and PIM are intended to support that distinction before reliance is established, and the gradient analysis shows that this support is graded rather than binary: a data substrate region with intermediate PBR carries intermediate model-directional reliability, and the framework’s measurements remain quantitatively meaningful across the full resolvability range.

A natural objection is that richer observation spaces — multiparametric imaging, genomics, longitudinal response patterns — may remove the shared-state condition. For the first case, this is correct in principle and is what the framework predicts (Proposition A6, Appendix): if richer inputs determine the contextual variable, the shared-state region disappears and PBR falls toward zero. For the second case, where observation is compatible with multiple generating laws at the same input, additional observation does not itself resolve the non-uniqueness. The results therefore do not preclude accurate state-to-law prediction; they define the conditions under which the observed state alone is insufficient, and distinguish those that richer observation can repair from those it cannot.

### Limitations

The Yale data substrate provides the imaging signal only, without the latent determinants *φ* that would resolve direction; the lesion proxy is an automated intensity-based summary rather than a structured expert annotation. This is deliberate: it ensures the empirical test matches the theoretical claim, which concerns what can and cannot be recovered from the observed state alone. The branching documented here is therefore a property of the observed state under a fixed law f; how an external regime change can itself alter f is a distinct failure mode outside the scope of this analysis.

Transitions may include multiple observations per subject; subject-level splitting mitigates but does not eliminate within-subject correlation. PBR is estimated from matched-state bins and is therefore dependent on a defensible binning choice; the sensitivity analysis across 16 modality–bin configurations (range 0.299–0.353; CV 4.3%) addresses this directly.

Deeper neural architectures might improve aggregate accuracy marginally but cannot resolve structural non-uniqueness when the input itself is compatible with multiple outcomes. The four-modality MLP experiment confirms this empirically: added capacity does not close the gap to majority-branch prediction in branching bins, and does not exceed the data substrate-determined ceiling in non-branching bins either.

## Conclusion

A state-only model can meet acceptable thresholds of aggregate accuracy while being structurally blind to trajectory direction in the data substrate regions where directional accuracy matters most. Standard validation may therefore certify models whose directional output, in these regions, does not exceed majority-branch prediction, while the same models recover direction with accuracy 0.96–1.00 in the complementary regions of the same data substrate where the observed state encodes direction. PBR and PIM are pre-reliance measures that identify this boundary before reliance is established, are computable without access to model weights, and index a continuous data substrate-resolvability gradient — captured by a near-deterministic Spearman correlation between minority-branch fraction and model directional accuracy across all four MRI sequences (*ρ* = −0.95 to −1.00) — rather than a binary partition.

## Data Sharing

The Yale Brain Metastases Longitudinal Data are publicly available through The Cancer Imaging Archive. All analysis code is available at a repository that can be requested from the author.

## Use of AI

Analyses were implemented by the author in Python (scikit-learn, numpy, pandas). A commercial large language model was used to review code for errors and to copy-edit manuscript prose; all scientific content, claims, analyses, and final text are the author’s own and were verified by the author.

## Contributors

MAE conceived the study, developed the formal framework, designed and conducted the analyses, and wrote the manuscript.

## 1 Appendix. Formal framework and proofs

This appendix establishes the formal framework underlying the main text. The central result, stated as Proposition A0 below, is that any deployed model — regardless of architecture, scale, or optimisation procedure — reduces to a single-valued predictor h : X → Z. All subsequent impossibility results follow from this reduction together with the assumption that the observed state x does not uniquely determine the contextual variable *φ* that governs trajectory direction. The fundamental claim is that state-only predictors cannot resolve subject-specific trajectory direction when the observed state is compatible with multiple admissible futures.

Formally, let X denote the space of observed states (for example, MRI features or lesion-burden summaries). Let Y denote the space of present observed states. Let ΔY denote the space of trajectory changes. Let Φ denote a space of contextual determinants not observable from x alone. The state map g : X → Y takes an observed state to its present-state label. The law of change f : X × Φ → ΔY maps observed state and context to trajectory change.

### 1.1 A0. The scalar-output constraint and reduction to h

Contemporary AI systems are trained by selecting parameters to minimise a loss function over observed data. The central structural fact is this: the loss *ℓ*(h(x; *θ*), z) must evaluate to a single scalar for each input–target pair (x, z). Suppose h could take two distinct values a≠ b at the same input x. *If ℓ(a, z) = ℓ(b, z) for all z, then h carries no information — the two outputs are indistinguishable under the loss. If ℓ(a, z)*≠ *ℓ(b, z) for some z, then the loss is not single-valued at x*, contradicting the requirement that *ℓ* evaluate to a single scalar for each (x, z) pair. In either case, h must be single-valued. This is not a limitation of current architectures that future architectures might overcome. It is a consequence of the optimisation framework itself.

Let h_j_(·; *θ*^k^) denote a predictor with architecture j and model scale k. For benchmark task B, target variable z, and observed sample {(x_n_, z_n_)}_n=1_^N^, the population objective is

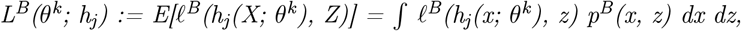

where p^B^ denotes the joint law induced by benchmark task B. Its empirical counterpart is

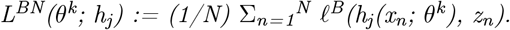

Training selects parameters by solving either the population problem *θ*^B^,_j_,^k^ ∈ argmin_*θ* L^B^(*θ*; h_j_), or the empirical problem *θ*^B^,_j_,^k^,^N^ ∈ argmin_*θ* L^BN^(*θ*; h_j_).

#### Proposition A0 (Reduction to h)

*Let Z be any target space. Consider any AI system trained on X for benchmark task B by minimising either L*^*B*^ *or L*^*BN*^ *over parameters θ, architecture j, or model scale k, and suppose the deployed system returns a determinate prediction for each input. Then:*

- the deployed system defines a single-valued predictor h : X → Z;
- whether the correct target is uniquely recoverable from x alone depends only on the information supplied in x, the admissible target values compatible with that information state, and the fact that h(x) is single-valued — not on the density p^B^, the expectation operator E[·], the architecture index j, the sample size N, the model scale k, or the optimiser;
- attention may therefore be restricted without loss of generality to h.

#### Remark.

If the system produces a distribution at inference time, the deployed system must still convert that distribution into a determinate prediction or decision. The above applies to any such deterministic selection rule.

***Proof***. Fix any benchmark task B, architecture j, model scale k, and trained parameter choice *θ*. The loss *ℓ*^B^(h(x; *θ*), z) must evaluate to a single scalar for each (x, z). If h could return two distinct values a≠ b at the same x, then either *ℓ*(a, z) = *ℓ*(b, z) for all z — in which case the two outputs are loss-equivalent and h carries no information distinguishing them — or *ℓ*(a, z)≠ *ℓ*(b, z) for some z, in which case the loss is not single-valued at that input, contradicting the scalar-output requirement. Therefore h(x; *θ*) is a single element of Z for each x, and the deployed system is a single-valued map h : X → Z. This proves (1). Whether the correct target is uniquely recoverable from x alone depends on two things: the admissible target values compatible with the information supplied in x, and the fact that h(x) is a single value. Neither depends on how *θ* was obtained. This proves (2), and (3) follows immediately. ■

#### Corollary A0.1 (Architectural irrelevance under contextual omission)

*Suppose* Δ*y = f(x, φ) with φ not determined by x. If there exists x* ∈ X exhibiting observational multiplicity — that is, *φ*_1_≠ *φ*_2_ with f(x, *φ*_*1*_)≠ *f(x, φ*_2_) — then no AI system trained only on X can uniquely recover subject-level direction of change at x. *If, in addition, f(x, φ*_1_) · f(x, *φ*_*2*_) ***<*** 0, then no such system can uniquely recover direction of change at x.

***Proof***. By Proposition A0, any such system reduces to a single-valued predictor h : X → Z. Since h(x*) is a single value and the two admissible outcomes are distinct, h(x*) cannot equal both. If the outcomes have opposite sign, h(x) cannot match both directions. ■

**Remark**. A natural objection is that with sufficient data the dominant regime at x will overwhelm the minority regime, so the model will correctly classify most cases. This is true only in a population-average sense. Direction of change here is a subject-level question: the individual subject is governed by one specific *φ*. If that subject lies on the minority branch, the population-optimal prediction is wrong for that subject. No increase in data volume can convert a single-valued predictor h(x) into a multi-valued one.

### 1.2 A1. Information limits

#### Theorem A1 (Information-theoretic non-uniqueness)

*Let X be an information space and Z a target space. Suppose there exists x* ∈ X and two distinct values z_1_≠ z_2_ ∈ Z such that both arise as correct target values under distinct admissible realisations sharing the same observed state x. *Then no single-valued predictor h* : *X* → *Z can uniquely determine the correct value at x*.

***Proof***. Since h is a single-valued map, h(x) is a single element of Z. Since z_1_≠ z_2_, no single element can serve as the correct value in both cases. ■

#### Remark.

The proof is short because the result is structural: it depends on no distributional assumptions, no regularity conditions, and no properties of the learning algorithm. Its force lies not in the proof technique but in the breadth of systems to which it applies.

#### Remark (Multimodal extension).

Theorem A1 applies regardless of the dimensionality or modality of x. If the observation space X is enlarged — for example by combining MRI, spectroscopy, and perfusion imaging into a composite observation — the result still holds provided two distinct latent states can produce the same composite observation. Enriching x enlarges the snapshot; it does not guarantee separation of the latent states that determine Δy.

### 1.3 A2. State does not imply direction of change

#### Theorem A2 (State–direction non-equivalence)

*Suppose g* : *X* → *Y is well-defined and there exists x* ∈ X exhibiting observational multiplicity. Then determination of y = g(x*) does not imply determination of* Δ*y from x* alone.

***Proof***. g(x*) is uniquely determined. But x* is compatible with distinct direction-of-change outcomes f(x, *φ*_*1*_)≠ *f(x, φ*_2_). By Theorem A1, no single-valued predictor on x alone can recover the correct outcome. ■

#### Theorem A3 (State-only impossibility)

*Let h* : *X* → Δ*Y be any predictor whose input is confined to x. If x* exhibits observational multiplicity, then h cannot represent both context-conditioned outcomes at x.

***Proof***. h(x*) is a single value. If h(x*) = f(x, *φ*_*1*_*) and h(x*) = f(x, *φ*_*2*_*), then f(x, φ*_1_) = f(x, *φ*_2_), contradicting observational multiplicity. ■

#### Corollary A3.1 (Structural error).

*Under the conditions of Theorem A3, every predictor h on x alone satisfies*

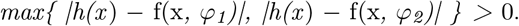

***Proof***. If both errors were zero, h(x) would equal both values, contradicting Theorem A3. ■

### 1.4 A3. Weighted loss minimisation under unresolved ambiguity

The preceding results establish that no single-valued predictor can eliminate error on both branches at a shared-state point. The next result characterises what such a predictor actually produces: the prediction is determined by loss geometry and relative branch prevalence, not by the true trajectory of the individual subject.

#### Proposition A2 (Weighted loss minimisation under unresolved ambiguity)

*Let hθ* : *X* → Δ*Y be a single-valued predictor and let ℓ* : Δ*Y* × Δ*Y* → *[0*, ∞*) satisfy ℓ(a, b) = 0 if and only if a = b. Suppose x*_*0*_ ∈ *X admits two distinct admissible targets* Δ*y*^*′*^≠ Δ*y*^*′′*^. *For positive weights w*^*′*^, *w*^*′′*^ ***>*** 0, define

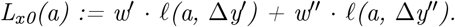

***Then:***

1. no a ∈ ΔY satisfies L_x0_(a) = 0;
2. any predictor minimising an objective with L_x0_ as its local contribution at x_0_ satisfies h*θ*(x_0_) ∈ argmin_{a ∈ ΔY} L_x0_(a);
3. the minimiser is determined by the loss geometry and relative weights, not by a uniquely determined true trajectory at x_0_.

***Proof***. Since Δy^*′*^≠ Δy^*′′*^, no a satisfies both a = Δy^*′*^ and a = Δy^*′′*^. Therefore at least one loss term is positive for every a, and since both weights are positive, L_x0_(a) > 0 for all a. This proves (1). The minimiser of the global objective must minimise L_x0_ locally at x_0_, proving (2). Since no action achieves zero loss, the selected action reflects loss geometry rather than a unique truth, proving (3). ■

#### Corollary A2.1 (Branch selection under 0–1 loss)

*If ℓ(a, z) = 1{a*≠ *z}, then L*_*x0*_(Δ*y*^*′*^*) = w*^*′′*^ *and L*_*x0*_(Δ*y*^*′′*^*) = w*^*′*^. *The minimiser is the branch with larger prevalence weight; w*^*′*^ ***>*** w^*′′*^ implies minimiser is Δy^*′*^, w^*′′*^ > w^*′*^ implies Δy^*′′*^; if w^*′*^ = w^*′′*^, both are minimisers.

***Proof***. Direct evaluation. ■

#### Corollary A2.2 (Weighted averaging under squared loss)

*If ℓ(a, z) = (a* − *z)*^*2*^, *the unique minimiser is*

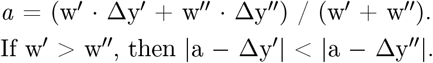

***Proof***. Differentiate L_x0_(a) = w^*′*^(a − Δy^*′*^)^2^ + w^*′′*^(a − Δy^*′′*^)^2^ and set equal to zero: (w^*′*^ + w^*′′*^) · a = w^*′*^ · Δy^*′*^ + w^*′′*^ · Δy^*′′*^. Strict convexity gives uniqueness. ■

#### Biological interpretation

Suppose at a given observed state, 70% of subjects are in the increasing regime and 30% are decreasing. Under squared loss, the optimal single-valued prediction is a weighted average of the two branches. This prediction is correct in sign for the 70% majority but wrong in sign for the 30% minority. The model confidently produces the prevalence-weighted compromise, which is the wrong answer for every minority-branch subject. In classification settings, the analogous effect is majority-branch selection rather than weighted averaging.

### 1.5 A4. Delayed observational determinacy

#### Theorem A5 (Delayed observational determinacy)

*Let x*_*0*_ *exhibit observational multiplicity, with admissible evolutions x*^*′*^*(t) and x*^*′′*^*(t) issuing from x*_*0*_ *cordecreasing to distinct directions* Δ*y*^*′≠*^ Δ*y*^*′′*^. *Suppose there exists t* **>** 0 such that x^*′*^(t)≠ *x*^*′′*^*(t*) and future direction is determinate on {x^*′*^(t*), x*^*′′*^*(t*)}. Then:

- future direction is not determinate at x_0_;
- later determinacy on {x^*′*^(t*), x*^*′′*^*(t*)} does not imply determinacy at x_0_;
- within this construction, determinacy arises only after the competing trajectories have separated in observed state space, since the sole source of directional information is the observed state, and at x_0_ it is shared.

***Proof***. Both Δy^*′*^ and Δy^*′′*^ are compatible with x_0_, so direction is not determinate there, proving (1). Later determinacy on separated states does not alter the shared-state ambiguity at x_0_, proving (2). At x_0_, any predictor receiving only x_0_ is single-valued and cannot distinguish the two cases, so determinacy requires separation, proving (3). ■

***Corollary A5.1***. *Later trajectory separation does not remove earlier ambiguity at x*_*0*_. *It shows only that the latent determinant of direction has become expressed in the observable state*.

***Proof***. Immediate from Theorem A5. ■

#### Biological interpretation

If direction of change is required at x_0_, waiting until t does not solve the problem; it delays resolution until the distinction has become expressed in the observable state. For downstream decisions that must be made in the window (t = 0, t), later observational clarity is not a substitute for earlier direction-of-change information.

### 1.6 A5. Context encoding does not resolve unobserved context

The preceding results establish that omitting *φ* from the predictor’s input creates an irrecoverable identifiability failure. A natural response is to include *φ* explicitly. This section establishes when doing so is well-posed, and why it does not circumvent the fundamental problem when *φ* is unobserved at deployment.

#### Proposition A6 (When contextual prediction is well-posed)

*If f(x, φ) is uniquely defined and locally supported by observed data for every (x, φ) in some domain D* ⊆ *X* × Φ, *then prediction by h* : *X* × Φ → Δ*Y is well-posed on D*.

***Proof***. The target map is single-valued and defined on D. The prediction problem reduces to approximation of a well-defined function. ■

#### Theorem A6 (Context encoding does not resolve unobserved context)

*Suppose x* exhibits observational multiplicity. Even if the predictor is instantiated as h(x, *φ*; *θ*) : X × Φ → ΔY, if the correct *φ* is not observed at deployment, the system must supply some 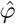. The prediction 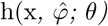 *is a single value and cannot equal both f(x, φ*_1_) and f(x, *φ*_*2*_*). In particular:*

- if 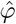 is fixed or defaulted, the predictor reduces to a function of x alone and Theorem A3 applies;
- if 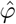 is estimated from x, the composite 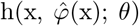 is single-valued in x and Theorem A3 applies;
- if X is enriched so that some proxy z(x) ≈ *φ* is available, h(x, z(x); *θ*) remains single-valued in x, and whenever z(x) takes the same value under both *φ*_1_ and *φ*_2_, multiplicity is unresolved.

***Proof***. The scalar-output constraint requires 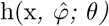 *to be a single element of* Δ*Y. In case (i), h depends only on x. In case (ii), the composite is a function of x alone; it is single-valued. In case (iii), z(x*) is the same regardless of the true *φ*, so h(x, *z(x*); *θ*) is again a single value. In all cases, the predictor cannot represent both admissible outcomes. ■

#### Remark.

The identifiability failure cannot be circumvented by architectural means unless the correct *φ* is genuinely observed and supplied at deployment. This is consistent with empirical observations that longitudinal imaging approaches improve predictive performance — not because multi-scan data inherently contains more information, but because serial imaging can make the contextual determinant *φ* observable through the trajectory itself.

### 1.7 A6. Local support failure and directionality degradation

#### Theorem A7 (Observational multiplicity under local support failure)

*Let x* ∈ X and *φ*_1≠_ *φ*_2_. If data are observed near (x, *φ*_*1*_*) but not near (x, φ*_2_), then f(x, *φ*_*2*_*) is not determined by the observed data*.

***Proof***. In the absence of observations near (x, *φ*_*2*_*), one may choose two admissible laws f and f that agree on the observed region and differ at (x, φ*_2_). Both are consistent with the data, but they disagree on the value in question. Hence that value is not determined. ■

#### Corollary A7.1 (Directionality degradation under imbalanced coverage)

*If one contextual regime is underrepresented near x*, the sign of f(x, *φ) under the underrepresented regime is not constrained by observations from the dominant regime. If PBR(x*) **>** 0, a predictor trained predominantly under one regime may assign the wrong direction of change under the other.

***Proof***. By Theorem A7, f(x, *φ*_*2*_*) is not determined by data near (x, φ*_1_). A predictor dominated by one regime has no observational basis for recovering the other, and may assign a value — including one with incorrect sign — reflecting the dominant regime. ■

### 1.8 A7. Definitions of PBR and PIM

#### Definition A1 (Predictive Blindness Risk)

*For x* ∈ *X, define*

*PBR(x) := sup {* |*f(x, φ*_*1*_) − *f(x, φ*_*2*_)| : *φ*_*1*_, *φ*_*2*_ ∈ Φ *}*.

PBR(x) = 0 if and only if direction of change at x does not depend on context.

#### Definition A2 (Prediction Indeterminacy Measure)

*For x* ∈ *X and φ*_*1*_, *φ*_*2*_ ∈ Φ *such that max{*|*f(x, φ*_*1*_)|, |*f(x, φ*_*2*_)|*}* ***>*** 0, define

*PIM(x; φ*_*1*_, *φ*_*2*_*) :=* |*f(x, φ*_*1*_) − *f(x, φ*_*2*_)| */ max{*|*f(x, φ*_*1*_)|, |*f(x, φ*_*2*_)|*}*.

PIM(x; *φ*_1_, *φ*_2_) > 0 if and only if the two contextual conditions produce distinct direction-of-change values at x. When PIM > 1, they produce values of opposite sign.

#### Proposition A4 (Vanishing of PBR)

*PBR(x) = 0 if and only if f(x, φ) is constant in φ*.

***Proof***. PBR(x) = 0 means |f(x, *φ*_1_) − f(x, *φ*_2_)| = 0 for all *φ*_1_, *φ*_2_, i.e., f is constant in *φ*. The converse is immediate. ■

#### Proposition A4.1 (PIM and sign reversal)

*If PIM(x; φ*_*1*_, *φ*_*2*_) ***>*** 1, then f(x, *φ*_1_) and f(x, *φ*_2_) have opposite sign.

***Proof***. Write M := max{|f(x, *φ*_1_)|, |f(x, *φ*_2_)|}. PIM > 1 gives |f(x, *φ*_1_) − f(x, *φ*_2_)| > M. If both values were of the same sign, the absolute difference would be at most M, contradicting the inequality. ■

